## SupplementaryData for "Neuropeptides from a praying mantis: What the loss of pyrokinins and tryptopyrokinins suggests about the endocrine functions of these peptides"

Jan A. Veenstra[1],

[1] INCIA, UMR 5287 CNRS, Université de Bordeaux, France

Content

| | |
|---|---|
| Fig. S1. Conceptual translation and predicted cleavage of neuroeptide precursors | 2 |
| Fig. S2. EFLamide receptors | 9 |
| Fig. S3. Ilp genes on chromosome 1 | 10 |
| Fig. S4. *Tenodera* sirps | 11 |
| Table S1. Mantodea genome SRAs analyzed for spots coding tryptopyrokinin and receptors | 12 |
| Table S2. Mantodea transcriptome SRAs analyzed for spots coding tryptopyrokinin and receptors | 13 |

**Fig. S1. Conceptually translated Tenodera sinensis neuropeptide precursors.** Predicted signal peptides are in yellow, putative convertase cleavage sites in red, predicted C-terminal amides in magenta and Cysteine residues in black. Parts of the precursor that are expected to end up in the mature peptide are in blueish grey. Note that since the processing sites of the three neuropeptide-like precursors (NPLPs) are difficult to predict, plausible basic amino acid residues have been indicate that might be important for cleavage.

**AKH**
MAISRVLVMLAMAAITLLAFCEAQVNFTPNWGKRTVLQDGPCKVSTDALMYIYKFIQSEAQKLVDCEKFGGN*

**ACP**
MARRFSERTLCWLLVALAVLATTWGQVTFSRDWNPGKRSQSEPPCNAAFRSAEEVCKVLVDEFRQIAACEIRSLLRY
QKEFEEKHADMFLERPEGR*

**Agatoxin-like**
MRTYLVVLACALLLLGQFVLPTVAGPYLEDADEGLEDYSDNNLERLLQDPAQKRSSLIYLFRRACIRRGGNCDHRPK
DCCYNSSCRCNLWGSNCRCQRMGLFQKWGK*

**Agatoxin-like**, alternative trancript
MRTYLVVLACALLLLGQFVLPTVAGPYLEDADEGLEDYSDNNLERLLQDPAQKRACIRRGGNCDHRPKDCCYNSSCR
CNLWGSNCRCQRMGLFQKWGK*

**Allatostatin-A**
MTRYLCYLPLLFLHLTHAVSPTPARGAPEEAGSGVVSSGSGSTAGLLLSHLEDPTTDGDNSDLDLMKRLYDFGLGKR
AYSYVSEYKRLPVYNFGLGKRSNKMYGFGLGKRSPASSQMYSFGLGKRDYDDYYQAEDEDEDIVDEEAGAEEDMQDD
DNIDLVDKRAGHMYSFGLGKRARPYSFGLGKRAPQQKMYGFGLGKRSPASSQMYSFGLGKRDYDDYYQAEDEDEDIV
DEEAGAEEDMQDDDNIDLVDKRAGHMYSFGLGKRARPYSFGLGKRAPQQKMYGFGLGKRGGSMYSFGLGKRGEGRLY
SFGLGKRPVGYGGRQSGSRFNFGLGKRSENSDLEDEMEQEAMEKRVVPKAGQRFAFGLGKREVAPGELEAVRMEENK
KNETHHSGNIVKRSLQYAFGLGKRTPSYSDMDDVEEFSRLVRRPYSFGLGKRVPMYNFGIGKRSEH*

**Allatostatin-B**
MQSGRGALLGFLLMVIVKSSCQEDPLPDQSRSQSANGEQTQDVTGVQVDEDKRAWHDLQGGWGKRGWQDLSGGWGKR
GWQDLSGGWGKRGWQDLNGGWGKRGWQDLNGGWGKRGWQDLNGGWGKRGWQDLNGGWGKRGWQDLNGGWGKRGWQDL
NGGWGKRGWQDLNGGWGKRGWQDLNGGWGKRGWQDLQGGWGKRDMDLEDDDIESEEDDEAVADEVKRAWSSLRAGWG
KRAADWANFRGSWGKRDPGWNNLKGLWGKRGDNWSRLSAAWGKRDS*

**Allatostatin-CC**
MGIHHHHQQHSCLLYSVLFLLVVCCSSTSLTERHSRSATAASLGADYPDYQSGVRYDEYPVVVPKRTAMLLDRIMVA
LQKAVDEDKSGRTYPSELAEKMDLQRRGQQKGRVYWRCYFNAVTCFKRK*

**Allatostatin-CCC**
MTSCTKLVLVLMLAVLGIVVQGKVLAQAGDKDRLLTEPDLVDDDGGIETALINYLFAKQMVNRLRNQMDISDLQRKR
SYWRQCAFNAVSCFGK*

**Allatotropin**
MRCSLSMCALLAVTLVAVLAMCGAASSSGSGSRGKPRTIRGFKNVALSTARGFGKRDGSADLVDQASPLDRFTESLP
VEWFVEEMRTNPELARIIVHKFVDADQDGELTAEELLRPVY*

**Baratin**
MATALCLMVLVVTLAAALPNTLLEDAQQAAHSASGIGGGRREELPTAIPLTSRAKSIHQQHLQGIGMGKPSSHNPHY
SAGQHMQYQKENLAHQQEGKTVSQYEKGYQYGVGKAALDKHVENALLKSELYGDPAAVNQYRYYGGANERKSNQQIT

YPSPTKRSFNPMMSYVLPSDELTNSRSRLKRELGFDPEDVLTVLSLWEADHHTNSESHQGINPSMYSYYGVQPQDIY
RPMEEEEEEAPEDEQEELGLDNDDTSQTDGDWLDSPVVQPSVYPHQFRLDRRGGYFYPQTYPQQHAQYQHPSHKREG
SHWGGFAKDKRFMVSRKRQAKADEDIPILTHLLNHPYHDAGLPINRRVVL*

**Bursicon-A**

MNCILTALFLLPLTYGVQQVTAADECQVTPVIHVLQYPGCVPKPIPSFACTGRCSSYLQVSGSKIWQMERSCMCCQE
SGEREASVSLFCPKAKPGERKFRKVTTKAPLECMCRPCTTVEESAVIPQEIAGYADEGPLSNHFRKSL*

**Bursicon-B**

MICKTQRVYMWFLTNLLFIVLPQLVICGDEDPACETLPSEIHLIKEEFDELGRLQRTCNGEVGVNKCEGACNSQVQP
SVITPTGFLKECYCCRESFLRERIITLTHCYDPDGMRLTQEGQATMDIKIREPADCKCFKCGDFSR*

**Calctionin, transcript A1**

MEWRRRAVTLLACLLVVVARTSSREIAQDIMDSHMRHLQENRRTVQLLKNLLEDLDVNMETVQKRTACYISAGMGHS
CDYRDIIGAADEAKYWKQEFIPGKRRRRDNF*

**Calctionin, transcript A2**

MEWRRRAVTLLACLLVVVARTSSREIAQDIMDRKRTACYISAGMGHSCDYRDIIGAADEAKYWKQEFIPGKRRRRDN
F*

**Calctionin, transcript B**

MEWRRRAVTLLACLLVVVARTSSREIAQDIMDSKTVERMKKWCANTGSDSCGNMYVPGSGDDEDYLNHGDNPGKRAL
LKRLSAYRNWARLLKCSTNTGDDSCGNGYVPGSGDDDDYLNNGNNPGKRALNVRPARCSTNTGDDSCGNGYVPGSGN
DNDYLNGGDNPGKRASPKLFVLHRLRCSTNTGDDSCGNGYVPGSGDDNDYLNGGGNPGKRAIPNLFGVLQHRPTRCS
TNTGDDSCGNGYVPGSGDDNDYLNGGGNPGKRASHSINFPEDGRYPQASSVKYASLKKFCDVFPMNPACRNLK*

**Carausius NPLP**

MAATPLLLVLLAALGLGTSHPNVEDNMIPNDEEILRALLQEGKSPRQQGQPEVEDSLGLPSDEESYNNLKAMMSLGA
TRPGSSGTHHFVSSSFPEIDARGFHESVFDGFGDYYPTWNRQKRDPLGINSRGFHDDVFSQDFGTFHTVKRDHKDDG
KTTILQRRKRDTTSEKQAGKKSQDQDDVGDEGTSADKRRPEMDGSGFHGDTFSGGFGDFWTMKKRRLGGSSFHGLDT
FSSGFGDFDTMKKRKPEMGASGFHGDTFSNGFGDFWTMKKDGQHPHLEHKRRPEMDSSGFHGDTFNSGFGDFWTMKK
RDYYGDTFNPGYGDLWSMKKRRPEMDSSGFHGDTFNSGFGDFWTMKKRRPEMDSSGFHGDTFNSGFGDFWTMKKRRP
EMDSSGFHGDTFNSGFGDFWTMKKRRPEMDSSGFHGDTFNSGFGDFWTMKKRRPEMDSSGFHGDTFNSGFGDFWTMK
KRKPEMDSSGFHGDLFNNGFGDFWTMKKRRPEMDSSGFYGDTFGGGFGDFWTMKKRRPEMDSSGFHGDTFNSGFGDF
WPMKKRSTSSKSHSKQATNTQH*

**Crustacean cardiactive peptide (CCAP),**

MQLKHVIMACTVVVLLALCGLPLASCDDVIVQKREVDPADMERLLDPKRKRPFCNAFTGCGKKRSDESMGTLVELNS
EPAVEELSRQILSEAKLWEAIQEARAEILRRRQDQASLVSLTFHFISNVKLQSK*

**CCHamide-1**, incomplete

????...SCLSYGHSCWGGHGKRSSSDIQTDIDVKDTQLFSSRAVPPLENQWELQDLSDIHLMQQMDNESRLSVKEQ
ENEDEVPDDLLFIADDNIGLPHKLARRRILYRPERKLEKMKILQTSP*

**CCHamide-2**

MGCLTASSWVTLALLIVILIVLQTDPCVAKRGCSAFGHSCFGGHGKRSEDEALAQPGIEQELTLQQDEIPATSQLQA
RMGLSTFLRQWLQSYRRTAGELDEK*

**CCRFamide**

MKKASWVVPAVVLLVLLVMLCGGNTEVDCSLSELRCELMCQLTELTRQCNKCRSRAPVRFGKRSGHEHHHYPPLLPP
PPPLPLDSEGQPVMPYEEKLKICCGQLLEFLLKSTAVAQRK*

**CNMamide**
MSCWTPLLWTLVTIAALCFEAEAAPEALHRRAGDPNIMPINALRESEELSDQEKMLREDVAALMEYLQQYQQQQQQ
GGEDEQVALDNGQENYPPVQQLPPALQVRLLRLQDGINKDDSKRGSYMTMCHFKICNMGRKRNLRYPWLRR*

**Corazonin**
MKSMHLRVLMICCVASVALAQTFQYSRGWTNGKKRSMEENPCNKLQAMRWLMTHCPFQFYLPHADVPKSGLPEASEE
PSLDFVERLRRLIPQEPSRR*

**CRF-like diuretic hormone**
MLTTAAILVLATFVTCGTAYYGGSPVLEAMAEPVSDYQTTSYLLPRLVAKYRAPHPSQGDWESASDPRFYLLTELDR
DASQAATRRVKRTGGGPSLSIVNPLDVLRQRLLLEIARRRMRQSQDQIQANRDFLQSIGKREVANRTDNEIADYIVS
PAVEKSGVPAPETNR*

**DH31**
MSCHNILLATALLIGAILMLSVTHAAESVPIASHRNNYITDLDADADSEYVLEMLTRLGQTIMRANDLENSKRGLDL
GLSRGFSGSQAAKHLMGLAAANYAGGPGRRRRSPDDIA*

**Ecdysis triggering hormone**
MAESSMCFWSLRHCLYLVSLLIVLYASEANGDEGPFFLKASKNVPRIGRRSEYDNFFLKASKNVPRIGRRREMAPLT
EGRDWGVWPWFKTADSIPGPSRRSDYYIHEEGETQPISWTTVEKTMEEAPELWKPELWRKMAEEMSGTDTKRGNNEQ
SQA*

**Eclosion hormone-1**
MEGRKTSALLILAMAAVVLLCVLEQAEGNGLGICIRNCAQCKKMLGPYFEGQLCAEACVKFKGKMIPDCEDVSSIAP
FLNKLE*

**Eclosion hormone-2**
MYHRTTMYLLLLTMMWCYGICSKEAITVCITNCGQCKQMFGSYFRGPVCAESCIASKGRLLPDCNNPNTLIGFLKRL
C*

**EFLamide**, last exon only.
..NGEKRFSLINKDNDAIFVKDSDPQRRSISSSGEVKSPSPRSLGSELLGKRYLHGGKHFKQEYELVKKYVVLSEVL
HKLSLLLNYFE*

**Elevenin**
MSRGCLRSISLPVILLSTVLLYHLAASEPKSVNCRRWVFHPTCRGVAAKRAYSQDSPAIFLVDNDRKGDSNLEEVLG
LYATPQPHSQSQNRAEQQRSNLRMQQPLAPWSAHGDGFKGESVYDWYLGRKRSRDNDVVYDY*

**FMRFamide**
MLRIVLALVLAAIASTYPTDNPISESPNILLATSDESDAPRDDSSNDALLNTLAAAMEDCESESEEESDEASKTSDE
DMLSVLPIRRCPSRNFLRFNRGRPDNFIRFGRGGREDSNFIRFGRGKSDSNFIRLGRGGGDRGNDNFIRFGRARSSN
FVRFGRSRPDNFIRFGRGRDSNFLRFGRSGSLEEPAPFGSSLQVDEDNNRVGRGGKAGSNFIRLGRAGSSSFIRLGR
DGEDQDEDIQERETRGRNTANFVRFGRRANNGNFLRFGRSSNSGELRRGKLTDRNFIRLGRSGSNMYEDDTNSGPAR
VERSENSRGFIRFGKRRDEEEDEDNKLVRLGRDVEEIQEPPVLPSSDANEDNDNNTSRTRRSIPYPKPEDEAQEGSY
PVIIATSSGGNSGHDDKVDTSSFRYYSPLIPNYILAPELSLLAPLSGAESTTKRARGDGHNRNYIRLG*

**Gonadulin**
MKTPVFLVSCPLFFAVMVHFTSGMPSEEDSCLRIISRIVIDDCSKWKRSIMLQEVGSVLRQHRSDIHSRSGHFSNKG
PLGELLGVPSHWVDDDLADVRRQYRQTIQHLWAECCSNKKKCSGDMFKGLCK*

**Glycoprotein hormone A2**
MVPVSWRLQCCSLLLVLVMLLSLVSRNSAREMDVWKRPGCHKVGHTRKISIPDCVEFHITTNACRGYCESWSVPSAI
DTLRVNPHQAITSVGQCCNIMDTEDVEVKVMCLDGARDLIFKSAKSCSCYHCKKD*

**Glycoprotein hormone B5**
MIPLNNNNNNQTFGPTTLVLLLVVLATLLGQTSAAMTMQENTLSNTLECHRRLYGYKVSKTDSAGRVCWDVISVMS
CWGRCDSNEISDWRFPYKRSHHPVCLHDRREVKYVDLRNCDEGAEPGTERYEYLEAISCRCMICKSSEASCEGLRYR
GQRSGPFLGGGR*

**Insulin-like growth factor (IGF)**, short transcript
MCSPQMWRSWTVLLVVVTALDLVRGTPLGRRQLCGRELADTLSSICFGRGYNDPFSASPGTEVPMYARSRTTRGVAD
ECCKTGCTWSTLEQYCNPRPPETSRSPQNVLTAEKHTSILRNTLESSLEASSSSSSSSSSSSKRRSSPRMSKKKKDR
KHDPSGKVEVAENVEQNANKIPPVIGTISPAYMRVPIVLLKRKAQDTAQN*

**Insulin-like growth factor (IGF)**, long transcript
MCSPQMWRSWTVLLVVVTALDLVRGTPLGRRQLCGRELADTLSSICFGRGYNDPFSASPGTEVPMYARSRTTRGVAD
ECCKTGCTWSTLEQYCNPRPPETSRSPQNVLTAEKHTSILRNTLESSLEASSSSSSSSSSSSKRRSSPRMSKKKKDR
KHDPSGKVRGHHKKKGRRGNNRCRCRRRRRRGKVEVAENVEQNANKIPPVIGTISPAYMRVPIVLLKRKAQDTAQN*

**short IGF-related peptide 1**
MKMWKLLVRLMALAAVCLCADVHGELTLMKRDIPQRYCGSNLVNVLQLVCRGNIYVVPDDKRSGSLLHNTVLPEEDD
ANLWRMEERRFPFRSRLSASSLVPRSFRRSKRQGVVQECCYYKGCTLSELSSYCGRR*

**short IGF-related peptide 2**
MWRTWILCLFTLLLVEAQLDKRSSTAKRYCGRNLIHILQLVCDSNYYSNTSPTIFNQKKSFPDDVWLQILEDNPVGD
DEPEFPFRSKPRASSFAHRVFRRHSRSRGVVDECCYDKGCTINELRGYCGSSR*

**short IGF-related peptide 3**
MKNFYIFCVAAIFCGALPIFTNVEAESLNSADSVVMGNPIETYCGSSLYIKLESVCNGNFNNKFHQEVTKCKLEPWG
IPCTIDSACCQTPCTERYIAGYCASTF*

**short IGF-related peptide 4**
MKKFDFLCFVAIFCGAFPIFTNVEADSFHVYPWEKRGDSPRKYCGNFLADILHLVCKGNYYSITGHNAENLSTQKKT
SGENEDSYWLQSLEEPSQEEFPFRSRLNSASMIRHRMFRRNAGSPGIVQECCVKGCTFSELSLYCAFR*

**short IGF-related peptide 5**
MWSAYIRLVALAALCLCTLAQAQSDLFQIGEKRNTPQKYCGRNLADILHLVCNGFYYPMFKKSADLDYDMNDAYWVE
SAPSPPQEQLPFPYRSRASATTVVNGGFRRRMRGIYDECCRKSCTILELSSYCGKR*

**invertebrate Parathyroid hormone**
MRTRLVFACSALVVLALLVIIPQTQGRPYRQKRVSDQRLAELETIMALRRMAGKLVSVPVGFGQVDPAKIGRRRRRS
AELLLQELLNAQANDVDDAEDSEAEDDIRELVGPHRPSQPWLSEWNRRVQVSIIYLL*

**Ion transport peptide**, transcript a
MAKQDNSIATLAKRALVCCLVVSVTTACLVRASPTSKLVIGHPLSKRSFFDIQCKGVYDKSIFARLDRICEDCYNLF
REPQLHSLCRKNCFTTEYFKGCLEALLLQDETEQIQTWIKQLHGAEPGV*

**Ion transport peptide**, transcript b
MAKQDNSIATLAKRALVCCLVVSVTTACLVRASPTSKLVIGHPLSKRSFFDIQCKGVYDKSIFARLDRICEDCYNLF
REPQLHSLCRSKCFSSRYFKGCLEALLLTEEEEKFSQMVDFLGKK*

**Leucokinin**
MWLATRGMNLILLLTAALATEEFLFPSRLPAIQEGLQTRLCQTAGVPYCSYNSPGYSDDQIPEPTEVKISTYTSGTG
KPMLLIAKNTDDLIRSENEADEAEPDAGVGELNPVKKDSAFSSWGGKRNENWNDKHNSEDPVVQLIKKKASAFSSWG
GKRASLDADPSEESDDYELFLPVVEPEDEENHVLKKSFSSWGGKRTFSNWGGKRVSNLDKPRRAFSSWGGKRAFSTW
GGKRDPSLHEAFGEAENAEKRTFASWGGKRKFSSWGGKRNALESIDKKAFSSWGGKRQLPCTNCNTVLAPDSYTGSA
NDKNLSIISFSMEDNGQTPEIFTPNKEKDYSRDIHQQNVFQNEKEESELLKALELGATSASPLKDANINTFLQDNMH
HMKKRDGRFSSITKHLTYPSIVVRGRFSSWGGKRAVKPLANRLSKTASPQTLERQYRRGGEFYAWGG*

**Myosuppressin**
MRNSCMMLIGVLAVVLVACVTAIPPPQCTSNNLEEIPPRVRKFCAALSTIYELSNAMETYLDDRVVRENTPMVDALP
KRQDVDHVFLRFGRRR*

**Natalisin**
MRPHAFLIIVVAWNHVVHGEETNPSLQEANHENATNSSVIAEKRVARSDLGGQGESEENPPPFWANRGRSLNLNGER
LRRHEPFFVEEPEWLVVEEEAPIPEEEDHHGECEESRKRRTSRLTGVDEAFWPSRGRRSGYKRPTGHDENAGRGAQD
LAGIVKMFHSTKGNLFNAKDSIKSHGEHRRLTNTPSIEEPFWAARGRSLFLEPRNRRSGLMESLEEPFWAARGRRSG
NSARGRRSEAFSGEEPFWAARGRRSDDPRGKRPESFSGEEPFWAARGRRFDGSQKVRRSQTSSGEEPFWAARGRRME
AGLHDSVSGKTPSPSTHNRLAEAGNPLKQAENQISEIARGRRSPYQKDHYVQFSSSEEPFWAARGKRGLLESLSAEE
PFWAARGKKQYPQMNNWWPIQEAAAALAEDNSEDESFWTALENKLLARRSVYDIVPM*

**Neuroparsin**
MSVRCTSVTLGVAILAMLLIQRCEGGSLLCKPCMGNECNLEPAGNCEHGVERDYCGWKVCAKGPGEHCGGPSDLMGK
CGEGMICTCGKCSGCSLATLECFFSSDQLHCI*

**Neuropeptide F-1**, transcript a
MMQSSALCWLVILGCLVLPQLAWSKPTDPEQLAAMADTLKYLQELDRYYSQVARPRFGKRSELRTLPEQETAPEESS
ERMWRRFVSRR*

**Neuropeptide F-1**, transcript b
MMQSSALCWLVILGCLVLPQLAWSKPTDPEQLAAMADTLKYLQELDRYYSQVARPSPRSESGRMHELSKVERALKML
RLQELDRFYSQRTRPRFGKRSELRTLPEQETAPEESSERMWRRFVSRR*

**Neuropeptide F-2**
MQQSPVILAGVLACLCVVSPCWSDPMAIGNEIHSRPTRPKVFTSPDELKTYLEQLSNFYAIAGRPRFGKRLAEPAMF
NSLSGPSPAAAATAARNFHFRFPGAAPVPSGSVRSDVYQMLFPYDE*

**Neuropeptide-like precursor**
MSPSPSQLLLAIAVFVFIVSFHKALTEDAGTEDQDADDKRTIGFMARIGAVPIMGKRYVASLARNGELPFLVRKEWH
KKMHPVMSGGRNIAELLNPPAGKRYIGALARSGNLPFATKRSQNEDGALENENEDVETLLKEAIDAGHLWRIELGAL
REKLLDENSVYPLLDFYETLNANGMTENHDEEKRAFPAIEPLGSVPFGRKRSVEALARAGYLPQLKPPQESEEYGQR
DSSEASEELLKRSAAGLSRNGNLKGLFQEILEGKRGGVGSLARNGYLRIGLDHFSGKRGGIGSLARSGTLRQKKFDE
EDNDELEELMKELNYLENYDDIARGLFGQDFSASGSEKRNIGSLARARDFPFKGVIKRIPFDEEEIIQKRNLASVLR
NRFAQQQGKRNLGSFMRSYGSSFVPTKKEDYTELDEQYKRNIGSMAKNWLLPEHIKNSKREVGNTLLYDGKDCSPAG
ADNEGEKKESFLGPQNDVTFHHVHKKSTHSTASDAPSYEAKNSTVSESDNAEQKSKSRNKREAYYSAAAPSEEYPLP
VLQNSDLYDYEDMADLLSGEGAPKKRFLGRIPQMGRNKPRTNSHSGRRRPQSRNI*

**Orcokinin**, transcipt A
MSPVASCRTVMMLVAASLVLQLAYAVPTQNDGYREYHGPDGEDDNVAHRLDSIAGGAHLIRELERQGHVPRQARGGL
DSLSGITFGGNKRLDSLSGITFGNQKRNFDEIDRAGFNSFVKKNFDEIDRAGFDSFVKRNFDEIDRVGFGSFVKRNT
PLLLARLYDKENN*

**Orcokinin**, transcript B
MSPVASCRTVMMLVAASLVLQLAYAVPTQNDGYREYHGPDGEDDNVAHRLDSIAGDASKKLQEINKDRWDEDMMKEL
YLTRNENAHSRIDSIGGGNIVRNTGHRRIATRGLDSIGGGNIIGRSLGGGTRTNILPSLDYLGDNHVRELDSIGGGN
IVGRDENSLYPYESLRSVDPIGGGNIVRAVDSIGGGNIVRNLDSLGGGNFVRSLDPIGGGNIVRSIDTIGGGNIVRS
SNFARALDSIGGGNIVRSIDTIGGGNIVRSSNFARALDSIGGGNIVRSVDPIGGGNIVRGLDPIGGGNIVRSSNFAR
ALDSIGGGNIVRSLDPIGGGNIVRSLDPIGGGNIVRSLDPIGGGNIVRSLDPIGGGNIVRDLGSFEGRRYFPLKSKN
KGSSRGH*

**Periviscerokinin**
MIATAFCFAITMLVLVSKASGTEPVIKHKDRRRNSGLIAFPRIGRSDLDLQFSYPASDFVSRWITVYKRQGEKKRQT
LIPFPRIGRSDDVAEESPVMVVDGAALPRNTWQIDAHNRLINTEMPWALVTFKDYARELGEVEDEDPILGNDNAHYT
GPQEKQ*

**Periplaneta NPLP**
MELWVRLILSLLMLLSPIQCTDEHPSDSLKTAIEAVSRRQRDLASFDSGYYPSGGLVGPRDELAFVAAPRDFAGDGQ
PENIGYGYQKSISSPSGMFAPPSQLAPVEQGYSTKSKTLENLILDYLGDDLKPDDDAQEIYYPNADIKRSAFRERYQ
NGRLEAMKKRYMGSSFRERTHQGPNENIEQKRGMMTDALIRKMEEDEEDERRDRGDDRNSNSPRYLELLRTMWRKYR
NENPNDIEIEDVSDDDVGEMLNYLRESGAIDEEDVDGIKTEIKKRQHYGNDYDFHMHNAAMGGWGGQGYRKRWNQRL
DGDENQKSSFLYSLKFVSPAANHEAIESLREEDEMVPDEHDKDILRLAAAESNRRDPAAWWLPALERGEAPEELFEA
PSEEEYQRLMLAQQGEHHVLPNRKRMKSNYDVPDILLAPEKRYLYDTAVIRKRFPVTKRSSNYYTSPPLLHHKNFIN
SDISERRKKKDAMGTSITTTDPKVAQELNQLFSSSSSHSESPLPFTATTHAPSTENSTSPNTTTKHNGTSETKHSNS
ASKKSAEQPIAMSREEAPLEIRKKSINWSEYFGIDRRRKKSEESHPVDEEWLVNQYFNTLAHEKPSLFHVDNDFPHT
IMRKGAMMQPFDTRVFDTDIFARNVQHKLESKKNARESNEDSTIDNMDDKLQHIEDQIVNEAVKYTGAHEGTTDSRE
IQEVKDKVLARLAAAYSLEKMRQALAEFKTSLQAQRMSKYNPENRKVEEGDEKHKRVAVKKEKVEDAKDEKDKRREN
EEDDSEEFLNNPVVVQPMSEGYMGKHFEFFNISEEECPIVDGIFNTCLLMGDEVGDHANLLISICILHQICYLCGPE
IGFPSAAACDHFFASEAHTACRGDPGCQHAAHKGMTFIQRERELTDNNCWNTPCIAHYFLHFPAPLPVSASVR*

**Pigment dispersing factor**
MKHLAALLVILYLVRMSLTSPVQQYEDDRYPTADKELNAVSPRELANWLMQLILHKGEANICTHKRNSEIINSLLGL
PKVLNEAGRK*

**Proctolin**
MCSRHILLLALVLMALYAATEARYLPTRSQDDRLDRLRELLRDLLESEIERSNVNNYERRMMFKREVPQIAAEQQLA
PVVSA*

**Prothoracicotropic hormone**
MKAFTVIQAAVLYAVTCCLCTASASLSEELREQSEEGSSPGCIGFCKKSEWLNTLLGSATKGTMNHEANQSAIVLSK
RNQNSSGSSFREQEEASCACQSSMSLVDLGQRTYPRYVTSAVCSNSLCRGYGTPCQSIYYITHVLRSKKAQTNAWAH
LQDVGPDGEITEMAAVVDTNGYPYTLPGNLSRKWKLDAIRVVAACLCMN*

**Pyrokinin**, *Tenodera sinensis* lacks this gene.

**relaxin/dilp7**, incomplete N-terminus
???.MLLHVSTVTAICVLIELSDSTSTEQELEELFKTRSNEDWEQAWHQERHARCQDRLLRHLYWACEKDIYRLYRR
NSQDEEDQPPSEDNSRWPFLSVLEAQVFLRDRRGARRRRAATSSITDECCVRTVGCTWEEYAEYCPSNKRFRKFV*

**RYamide**, alternative version
MKCSSVMLTVTIVASLAVLTSSATQFYASGRYGKRAEMSGPMFWTGSRYGRSSGSGTVAAALPGGGRLGDTVEVAAR
NERFFGGRRYGKRAEMSGPMFWTGSRYGRSSGSGTVAAALPGGGRLGDTVEVAARNERFFGGSRYGKRGVDQDQAAI
ADSPRGVLAVEDEASQVTCLYTGVTNLYRCYNRKENSSSEESVNSERTK*

**short Neuropeptide F**

MQGFPTIKCCTIALCFLIVAAEFVAGAPSYSDYETGVRDLYELLLQKEALENRLQAQQALAGQTTHEVVRKANRSPS
LRLRFGRRADPLLAAAASPFMEHSAESGIAEN*

**SIFamide**

MQKSGVATCVLLLVVILLAEVAMAAYKKPPFNGSIFGKRGTVVEYDAAGRALSAMCEIASEACSAWFSQSENK*

**SMYamide**

MQMGKTMFTFVMMLLLVTLMAQTTSSHRRIPFSGSMYGKRGGDSYDSKIKSISTMCEVAADICTIWFPPTEN*

**Sulfakinin**

MCAFFALRMLLLLTVGVYLALQHCATAAPSTSEVSAAGTSVQRARVHSFPRVRARLVPLEPSSDLLSDFIIDDEFAD
FNKRQSTDGKREKEFDDYGHMRFGKREQFDDYGHMRFGRSLD*

**Tachykinin**

MTHWRISALILVTLVFTVALCTPEESPKRAPSGFLGVRGKKSDTSSSSAVSYDFAEKRAPAMGFQGVRGKKDNDLML
DFDTADKRAPAMGFQGVRGKKDYDTDLLDLDYFDKRAPAMGFMGMRGKKADLDDILGKRAPALGFQGMRGKKDDWND
ESDMYKRAPSSGFHGMRGKKDFDDYMNVYPGDKRMGFMGMRGKKKEFDEEDYEEALSNGDFWNEEEYLEGESKRAPA
AGFFGMRGKKGPSSGFFGMRGKKGPSAGFFAMRGKKAPGSGFMGMRGKKAPSAGFMGMRGKKDYEDEGDSLESLLQQ
LGYEQAKGREKRTSGQWSMDQGKTILILSIHRLST*

**Trissin**

MAGTTAHLIFLVTGLVLCTWSVALSCDSCGRECQAACGTRNFRTCCFNYLRKRSDGNALDRPGLRLELLVVPELAAR
YWEDHLKPLHPVPVFTEPEDTTENTRGHMQLIYNP*

**Tryptopyrokinin, *Tenodera sinensis* lacks this gene.**

**Vasopressin (Inotocin)**

MSQSEMKTQWFLMFVTTICISSACLITNCPKGGKRTPYDKQDTIKQCARCGPARLGHCYGPAICCGPQIGCLVATPD
TARCLEEAASPVPCVAPTGPVCGVGDTLGRCTANGVCCTHDSCSLDVSCRITVGDTLELMDGGPMFNLYNRRHSLID
SQ*

```
Locusta     MRPAGAAVALAALLPLLLAAGAAASDAAVAPPAAEPAYYSARYRLVGTLCQGVVLAVGLA    60
Tenodera    ----------------------MMNYSVESLEREDAEYYSYRYRTIGTIFQGIILIVGVL    38
                                    . :. :    :  *** *** :**: **::* **:

Locusta     GNLLVVAVVCGARSMRSPTNCYLVSLAVADCLVLVASVPNEIASYYLVGNQWLWGDAGCA    120
Tenodera    GNVMVVVVVHRTRSMRTPTNCYLVSLAVADCLVLVVTVPQEIASYYLVGNLWLWGKAGCN    98
            **::**.**  :****:******************.:**:********** ****.***

Locusta     AFVFSQNLGINASALSLVAFTVERYVAICRPLRSHALRSVARARRVSLLAWAAAAAYSAP    180
Tenodera    IYVFCQYLGINASSLSLVAFTVERYIAICKPLHAHAVCTVSRAQRIAFGVWIFATLYCSP    158
             :**.* ******:***********:***:**::**: :*:**:*::: .*  *: *.:*

Locusta     WLLLAATRPLRYRGLPELRACAFRLERARYLPYFLCDLLLFYAAPLLLCCVLYALIARAL    240
Tenodera    WILLSGTSPLLYKGLPDGESCGFKLPRAHYLAYFFTDLVLFYVIPLIVSCVLYSLIAKVL    218
            *:**:.* ** *:***: .:*.*:* **:** **: **:***. **::.****:***:.*

Locusta     FRRAALAASGGAGLSPHASAAGVDARCQVVRMLAAVVAAFAALWLPYRGLLVYNSFATLL    300
Tenodera    RSRRIAGS-------AQQIESTAPAREQVVKMLAVVVVVFATLWLPYRGMLVYNSFATLF    271
              *   .:        :   : . ** ***:***.**..**:*******:*********:

Locusta     SGDKYMDLWFLLFAKTCVFVNSAINPILYNAMSAKFRRAFRRALLRCTRRAAAAAAPADG    360
Tenodera    SGQRFMNLWFLMFAKTCVYINCAINPILYNAMSVKFRRAFRRTLCGGGISAKDS------    325
            **:::*:****:******::*.***********.********:*      *  :

Locusta     PLSGSGGTRLMV---------------------       372
Tenodera    -SSSRGSAGLLQQKKHPSAHRTHSLSTTLSTKR       357
              *. *.: *:
```

*Locusta*: QGT41395.1, coding sequence the *Tenodera* sequence as deduced from the genome:

*Tenodera* deduced coding sequence of ELLamide receptor.
ATGATGAACTATAGTGTGGAGTCGCTGGAGCGTGAAGATGCTGAATATTACTCTTATCGG
TACCGTACCATCGGTACCATCTTCCAGGGCATCATACTCATTGTTGGTGTACTGGGCAAC
GTCATGGTCGTCGTCGTGGTGCACCGTACACGTTCAATGCGCACTCCTACCAACTGTTAC
TTGGTGAGCCTAGCAGTGGCGGATTGCCTCGTGCTCGTGGTGACGGTGCCACAAGAGATA
GCCAGCTACTACCTGGTCGGCAACCTGTGGCTGTGGGGCAAGGCCGGCTGTAACATCTAC
GTGTTTTGTCAATATCTGGGGATCAACGCCTCCTCGCTGAGCCTGGTAGCCTTCACTGTG
GAGCGCTACATCGCCATCTGCAAACCTCTGCACGCTCATGCGGTGTGTACAGTGTCCAGG
GCCCAGCGTATTGCCTTCGGTGTCTGGATATTCGCCACCCTGTACTGCTCGCCCTGGATC
CTGCTCTCCGGCACAAGTCCTCTTCTCTATAAGGGTTTGCCAGACGGCGAGAGCTGTGGT
TTCAAGCTTCCTCGTGCACACTACCTGGCCTATTTCTTCACAGACCTCGTGCTTTTCTAC
GTAATACCTCTCATAGTATCCTGTGTGCTGTACTCTCTGATCGCCAAGGTCCTGCGCAGT
CGTCGAATTGCTGGATCTGCGCAACAAATTGAGTCTACTGCACCAGCCAGAGAACAGGTG
GTGAAGATGCTGGCAGTAGTAGTGGTGGTGTTCGCTACCTTATGGCTGCCATACAGAGGG
ATGCTGGTTTACAACTCCTTCGCTACCTTGTTCTCCGGACAGAGATTCATGAACCTTTGG
TTCCTCATGTTCGCCAAGACGTGTGTTTACATCAACTGTGCCATCAACCCAATCCTGTAC
AATGCTATGTCTGTCAAGTTCAGGAGAGCATTCCGGAGGACGCTGTGTGGAGGAGGAATT
TCAGCGAAAGACAGCAGCAGCAGTCGAGGATCAGCAGGCCTGTTACAGCAGAAGAAGCAC
CCTTCTGCACATCGCACACATTCCTTGTCTACAACACTCTCAACCAAGCGATAG

**Fig. S2**. EFLamide receptors. Sequence alignment of EFLamide receptors from *Locusta migratoria* and *Tenodera sinensis* and the coding sequence of the *Tenodera* receptor.

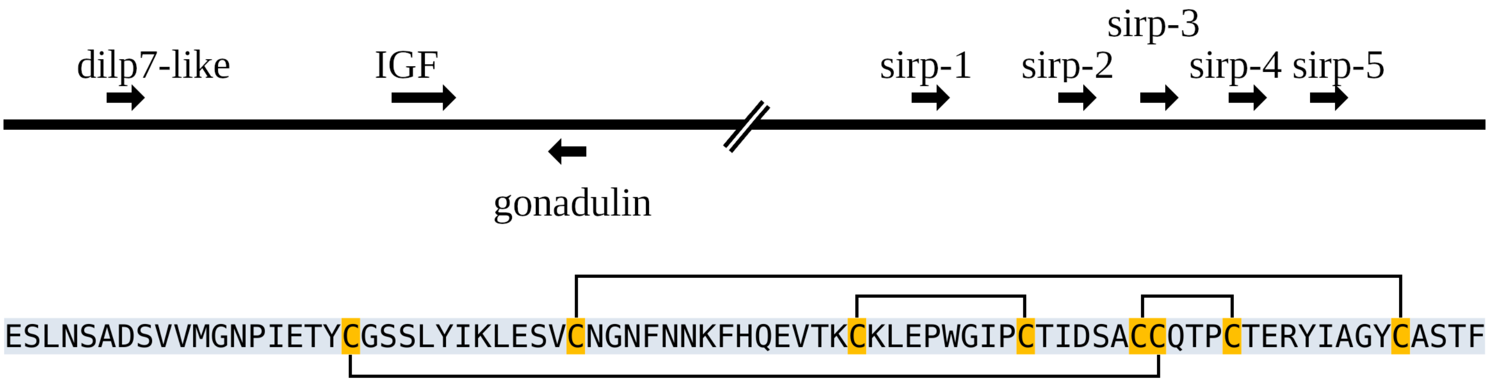


**Fig. S3.** Ilp genes on chromosome 1. Top schematic representation of the relative orientation of the various *Tenodera* ilp genes on chromosome 1. The dilp7 ortholog, IGF and gonadulin are present in a very similar configuration as in cockroaches. The five sirp genes are locate next to one another on a different fragment of the same chromosome. These two fragments are separated by more than 40,000,000 bp.

```
T.angustipennis-1      MKMWKLLVRLMALAAVCLCADVHGELTLMKRDIPQRYCGSNLVNVLQLVCRGNIYVVPDD     60
T.sinensis-1           MKMWKLLVRLMALAAVCLCADVHGELTLMKRDIPQRYCGSNLVNVLQLVCRGNIYVVPDD     60
                       ************************************************************

T.angustipennis-1      KRSGSLLHNTVLPEEDDANLWRMEERRFPFRSRLSASSLVPRSFRRSKRQGVVQECCYYK     120
T.sinensis-1           KRSGSLLHNTVLPEEDDANLWRMEERRFPFRSRLSASSLVPRSFRRSKRQGVVQECCYYK     120
                       ************************************************************

T.angustipennis-1      GCTLSELSSYCGRR    134
T.sinensis-1           GCTLSELSSYCGRR    134
                       **************


T.angustipennis-2      MANTKMWRTWILCLFTLLLVEAQLDKRSSTAKRYCGRNLIHILQLVCDSNYYSNTSPTIF     60
T.sinensis-2           -----MWRTWILCLFTLLLVEAQLDKRSSTAKRYCGRNLIHILQLVCDSNYYSNTSPTIF     55
                            *******************************************************

T.angustipennis-2      NQKKSFPDDVWLQILEDNPVSDDEPEFPFRSKPRASSFAHRVFRRHSRSRGVVDECCYDK     120
T.sinensis-2           NQKKSFPDDVWLQILEDNPVGDDEPEFPFRSKPRASSFAHRVFRRHSRSRGVVDECCYDK     115
                       ********************.***************************************

T.angustipennis-2      GCTINELRGYCGSSR   135
T.sinensis-2           GCTINELRGYCGSSR   130
                       ***************


T.angustipennis-3      MKNFYIFCVAAIFCGALPISTNVEAGRLNAYAREKRDVIISEYPKIYCGRELNEALQIVC     60
T.sinensis-3           MKNFYIFCVAAIFCGALPIFTNVEAESLNSA----DSVVMGNPIETYCGSSLYIKLESVC     56
                       ******************* *****  **:      .*::.:  : *** .*   *: **

T.angustipennis-3      DGKFNPIPVGGTAARGKRDYDIRRRPRGVVDECCKNPCTREVLEQYCA---   108
T.sinensis-3           NGNFNNKFHQEVTKC---KLEPWGIPCTIDSACCQTPCTERYIAGYCASTF   104
                       :*:**      .:     . :    *  : . **:.***.. :  ***

T.angustipennis-4      MKKFDFFCFVAIFCGAFPIFTNVEADSFHVYPWEKRGDSPRKYCGNFLADILHLVCKGNY     60
T.sinensis-4           MKKFDFLCFVAIFCGAFPIFTNVEADSFHVYPWEKRGDSPRKYCGNFLADILHLVCKGNY     60
                       ******:*****************************************************

T.angustipennis-4      YSITGHNAENLSTQKKTSGENEDSYWLQSLEEPSQEEFPFRSRLNSASMIRHRVFRRNAG     120
T.sinensis-4           YSITGHNAENLSTQKKTSGENEDSYWLQSLEEPSQEEFPFRSRLNSASMIRHRMFRRNAG     120
                       *****************************************************:******

T.angustipennis-4      SPGIVQECCIKGCTFSELSLYCAFR     145
T.sinensis-4           SPGIVQECCVKGCTFSELSLYCAFR     145
                       *********:***************


T.angustipennis-5      MWSAYIRLVALAALCLCMLAQAQSDLFQIGEKRNTPQKYCGRNLADILHLVCNGFYYPMF     60
T.sinensis-5           MWSAYIRLVALAALCLCTLAQAQSDLFQIGEKRNTPQKYCGRNLADILHLVCNGFYYPMF     60
                       ***************** ******************************************

T.angustipennis-5      KKSADLDYDMNDAYWVESAPSPPQEQLPFPYRSRASATTVVNGGFRRRMRGIYDECCRKS     120
T.sinensis-5           KKSADLDYDMNDAYWVESAPSPPQEQLPFPYRSRASATTVVNGGFRRRMRGIYDECCRKS     120
                       ************************************************************

T.angustipennis-5      CTILELSSYCGKR     133
T.sinensis-5           CTILELSSYCGKR     133
```

**Fig. S4.** *Tenodera* sirps. Sequence comparison of the five short IGF-related peptides from *Tenodera angustipennis* and *T. sinensis*. Note that most of them are well conserved, but that sirp-3 is very different. In *T. sinensis* it has 8 cysteine residues, but in *T. angustipennis* only 6.

| SRA | Species | TryptoPK | PeriVKR | TryptoPKR | PKRA | PKRB |
|---|---|---|---|---|---|---|
| SRR18233046 | *Amantis wuzhishana* | absent | present | absent | absent | absent |
| SRR18210554 | *Anaxarcha graminea* | absent | present | absent | absent | absent |
| SRR18218783 | *Anaxarcha sinensis* | absent | present | absent | absent | absent |
| SRR18217670 | *Anaxarcha sp.* | absent | present | absent | absent | absent |
| SRR18215879 | *Anaxarcha tianmushanensis* | absent | present | absent | absent | absent |
| SRR18246376 | *Gonypeta brunneri* | absent | present | absent | absent | absent |
| SRR18246425 | *Gonypeta sp.* | absent | present | absent | absent | absent |
| SRR15584233 | *Hierodula chinensis* | absent | present | absent | absent | absent |
| SRR18217671 | *Hierodula latipennis* | absent | present | absent | absent | absent |
| SRR16641550 | *Hierodula longa* | absent | present | absent | absent | absent |
| SRR16955306 | *Hierodula maculata* | absent | present | absent | absent | absent |
| SRR16955305 | *Hierodula sp.* | absent | present | absent | absent | absent |
| SRR15590770 | *Hierodula zhangi* | absent | present | absent | absent | absent |
| SRR18233043 | *Odontomantis sp.* | absent | present | absent | absent | absent |
| SRR18245581 | *Sinomiopteryx sp.* | absent | absent | absent | absent | absent |
| SRR18212399 | *Statilia agresta* | absent | present | absent | absent | absent |
| SRR16641552 | *Statilia flavobrunnea* | absent | present | absent | absent | absent |
| SRR16641551 | *Statilia maculata* | absent | present | absent | absent | absent |
| SRR16955304 | *Statilia sp.* | absent | present | absent | absent | absent |
| SRR16641549 | *Tenodera angustipennis* | absent | present | absent | absent | absent |
| SRR16641553 | *Tenodera aridifolia* | absent | present | absent | absent | absent |
| SRR18218046 | *Tenodera sp.* | absent | present | absent | absent | absent |
| SRR18218491 | *Titanodula formosana* | absent | present | absent | absent | absent |

| SRA | Species | TryptoPK | PeriVKR | TryptoPKR | PKRA | PKRB |
|---|---|---|---|---|---|---|
| SRR18233328 | *Acromantis hesione* | present | present | absent | absent | absent |
| SRR18210474 | *Acromantis japonica* | present | present | absent | absent | absent |
| SRR18217669 | *Arria brevifrons* | present | present | absent | absent | absent |
| SRR18245254 | *Arria pallida* | present | present | absent | absent | absent |
| SRR18233417 | *Arria pura* | present | present | absent | absent | absent |
| SRR18245238 | *Astyliasula major* | present | present | absent | absent | absent |
| SRR18246373 | *Caliris sp.* | present | present | absent | absent | absent |
| SRR18231900 | *Leptomantella sp.* | present | present | present | absent | absent |
| SRR18210593 | *Phyllothelys sinense* | present | present | absent | absent | absent |
| SRR18215878 | *Phyllothelys werneri* | present | present | absent | absent | absent |
| SRR18237807 | *Pseudempusa pinnapavonis* | present | present | absent | absent | absent |
| SRR18210333 | *Sinomiopteryx sp.* | present | present | absent | absent | absent |
| SRR18210332 | *Sinomiopteryx sp.* | present | present | absent | absent | absent |
| SRR18246465 | *Theopompa maculosa* | present | present | absent | absent | absent |
| SRR18246455 | *Theopompa maculosa* | present | present | absent | absent | absent |
| SRR18210254 | *Theopompa ophthalmica* | present | present | absent | absent | absent |
| SRR18218670 | *Theopropus sinecus* | present | present | absent | absent | absent |
| SRR18232936 | *Theopropus sp.* | present | present | absent | absent | absent |
| SRR18233413 | *Theopropus sp.* | present | present | absent | absent | absent |

**Table S1.** Genome SRAs from Mantodea. These SRAs were analyzed for the presence of spots containing coding sequences for tryptopyrokinin (TryptoPK), or orthologous to four *Periplaneta* GPCRs that are the receptors for periviscerokinin (PeriVKR), tryptopyrokinin, the pyrokinin-1 receptor (TryptoPKR), or the two other pyrokinin receptors (PKRA and PKRB). Note that in genome SRAs a GPCR is readily detected, there is only one SRA in which no spots for the periviscerokinin receptor were found. In orange the *Leptomantella* SRA that contains spots for the pyrokinin-1 receptor.

| SRA | Species | TryptoPK | PeriVKR | TryptoPKR | PKRA | PKRB |
|---|---|---|---|---|---|---|
| SRR2230505 | *Amorphoscelis pulchra* | absent | present | absent | absent | absent |
| SRR2230519 | *Chaeteessa sp.* | absent | present | absent | absent | absent |
| SRR2230520 | *Choeradodis rhombicollis* | absent | absent | absent | absent | absent |
| SRR2230539 | *Eremiaphila braueri* | absent | present | absent | absent | absent |
| SRR2230561 | *Hierodula patellifera* | absent | absent | absent | absent | absent |
| SRR1185954 | *Hymenopus coronatus* | absent | absent | absent | absent | absent |
| SRR1185955 | *Hymenopus coronatus* | absent | absent | absent | absent | absent |
| SRR11669710 | *Leptomantella albella* | absent | absent | absent | absent | absent |
| SRR921615 | *Mantis religiosa* | absent | absent | absent | absent | absent |
| SRR2230571 | *Mantoida sp.* | absent | present | absent | absent | absent |
| SRR2230587 | *Omomantis zebrata* | absent | absent | absent | absent | absent |
| SRR1811980 | *Orthodera novaezealandiae* | absent | present | absent | absent | absent |
| SRR2230593 | *Oxyopsis gracilis* | absent | present | absent | absent | absent |
| SRR2230598 | *Paraoxypilus sp.* | absent | present | absent | absent | absent |
| SRR2230605 | *Phasmomantis sumichrasti* | absent | absent | absent | absent | absent |
| SRR2230615 | *Pseudogalepsus nigricoxa* | absent | present | absent | absent | absent |
| SRR2230554 | *Pseudovates hofmanni* | absent | absent | absent | absent | absent |
| SRR2230621 | *Rhombodera basalis* | absent | absent | absent | absent | absent |
| SRR2230627 | *Sphodromantis lineola* | absent | absent | absent | absent | absent |
| SRR1811993 | *Stagmatoptera biocellata* | absent | present | absent | absent | absent |
| SRR2230636 | *Theopropus elegans* | absent | absent | absent | absent | absent |
| SRR2230638 | *Thesprotia graminis* | absent | present | absent | absent | absent |
| SRR11729950 | *Titanodula formosana* | absent | present | absent | absent | absent |

| SRA | Species | TryptoPK | PeriVKR | TryptoPKR | PKRA | PKRB |
|---|---|---|---|---|---|---|
| SRR2230497 | *Acanthops sp.* | present | present | absent | absent | absent |
| SRR1811954 | *Acontista multicolor* | present | present | absent | absent | absent |
| SRR2230500 | *Acromantis sp.* | present | absent | absent | absent | absent |
| SRR2230504 | *Ameles decolor* | present | absent | absent | absent | absent |
| SRR1811961 | *Brunneria borealis* | present | present | absent | absent | absent |
| SRR1811963 | *Cheddikulama straminea* | present | present | absent | absent | absent |
| SRR1811965 | *Creobroter pictipennis* | present | absent | absent | absent | absent |
| SRR2230526 | *Danuria thunbergi* | present | present | absent | absent | absent |
| SRR2230528 | *Deroplatys lobata* | present | absent | absent | absent | absent |
| SRR921590 | *Empusa pennata* | present | absent | absent | absent | absent |
| SRR2230536 | *Ephestiasula pictipes* | present | absent | absent | absent | absent |
| SRR2230540 | *Euchomenella sp.* | present | absent | absent | absent | absent |
| SRR2230549 | *Gonatista grisea* | present | present | absent | absent | absent |
| SRR2230556 | *Harpagomantis tricolor* | present | present | absent | absent | absent |
| SRR2230559 | *Heterochaeta occidentalis* | present | present | absent | absent | absent |
| SRR2230562 | *Humbertiella sp.* | present | present | absent | absent | absent |
| SRR2230563 | *Idolomantis diabolica* | present | absent | absent | absent | absent |
| SRR2230568 | *Liturgusa sp.* | present | present | absent | absent | absent |
| SRR921620 | *Metallyticus splendidus* | present | absent | present | absent | absent |
| SRR2230577 | *Miomantis binotata* | present | absent | absent | absent | absent |
| SRR2230584 | *Nilomantis floweri* | present | present | absent | absent | absent |
| SRR2230591 | *Orthoderella ornata* | present | present | absent | absent | absent |
| SRR2230594 | *Oxyothespis dumonti* | present | absent | absent | absent | absent |
| SRR2230601 | *Parasphendale sp.* | present | absent | absent | absent | absent |
| SRR1811985 | *Phyllocrania paradoxa* | present | absent | absent | absent | absent |
| SRR2230606 | *Phyllothelys werneri* | present | present | absent | absent | absent |
| SRR1811986 | *Popa spurca* | present | absent | absent | absent | absent |
| SRR2230614 | *Pseudempusa pinnapavonis* | present | present | absent | absent | absent |
| SRR2230624 | *Sibylla pretiosa* | present | absent | absent | absent | absent |
| SRR2230635 | *Telomantis lamperti* | present | present | absent | absent | absent |
| SRR2230641 | *Tropidomantis tenera* | present | present | absent | absent | absent |

**Table S2.** Transcriptome SRAs from Mantodea. For explanation see Table S1. Note that GPCRs are not easily detected as in genome SRAs, but that nevertheless in *Metallyticus splendidus* a spot for a pyrokinin-1 receptor was found.
